# Characterization of enterovirus D68 infection in four nonhuman primate species

**DOI:** 10.1101/2022.04.16.487524

**Authors:** Justin Dearing, Abigail Conte, Catherine Brooks, Anna Zimina, Rhiannon Rivas, Sharla M. Birch, Danielle R. Adney, Maggie Li, Adriana Rascon, George A. Belov, Andrew Pekosz, Meghan S. Vermillion

## Abstract

Human enterovirus D68 (EV-D68) is a globally reemerging respiratory pathogen that is associated with the development of acute flaccid myelitis (AFM) in children. Currently, there are no approved vaccines or treatments for EV-D68 infection, and there is a paucity of data related to the virus and host specific factors that predict disease severity and progression to the neurologic syndrome. Published animal models of EV-D68 infection to date have been limited to mice, cotton rat and ferrets, and investigation of the susceptibility of nonhuman primate (NHP) species to contemporary EV-D68 isolates has not yet been reported. In this study, we challenged juvenile NHPs – cynomolgus macaques, rhesus macaques, pigtailed macaques, and African green monkeys – with one of five different 2014 or 2018 EV-D68 isolates by the respiratory route. Animals were monitored for clinical respiratory and neurologic signs, and serially collected nasal swabs, bronchoalveolar lavage fluid (BALF) and cerebrospinal fluid (CSF) were evaluated for EV-D68 RNA and infectious virus. Infection with 2014 and 2018 EV-D68 isolates resulted in mild respiratory and gastrointestinal disease in some animals, but no evidence of neurological disease. Neither EV-D68 RNA nor infectious virus could be detected from any sample collected from animals challenged with 2014 EV-D68 isolates. Limited viral shedding – based on viral RNA quantified from nasal swabs and BALF – was detected from some animals infected with 2018 EV-D68 isolates. No virus was detectable in CSF. The rate of seroconversion was 100% for cynomolgus macaques infected with the 2018 EV-D68 isolates, but averaged between 0-50% for the 2014 isolates. Based on the results of this study, there is some evidence that infection with 2018 EV-D68 isolates may be more reliable at establishing limited infection than 2014 EV-D68 isolates. Regardless of virus isolate, however, EV-D68 infection of juvenile NHP species resulted in mild and nonspecific clinical disease and limited viral shedding. These data suggest that further refinements to the NHP model system (e.g., immunosuppression and/or direct viral inoculation) may be required to reproduce EV-D68 infection of the central nervous system and the associated AFM phenotype.

## Introduction

Human enterovirus D68 (EV-D68) is a non-polio enterovirus that can cause severe respiratory illness, and infection has been linked with a neurologic syndrome known as acute flaccid myelitis (AFM), which is most commonly reported in children (Lang et al., 2014). Over the past two decades, the incidence of reported EV-D68 infections has continued to increase worldwide, and periods of heightened detection have coincided with biennial outbreaks of AFM (Chong et al., 2018; Dyda et al., 2018; Greninger et al., 2015; Kamau et al., 2019; Knoester et al., 2019; Messacar et al., 2016). At present, there are no approved vaccines or therapies for EV-D68 infection, and the existing treatments for AFM patients are non-specific and largely anecdotal. Development and testing of more targeted EV-D68 vaccines and therapies relies on a comprehensive understanding of EV-D68 pathogenesis *in vivo*, including cell receptor usage, mechanisms of viral dissemination and tissue tropisms, as well as host innate and adaptive immune responses.

Experimental EV-D68 infection has been described in mice, cotton rats, and ferrets. Productive infection of mice with either historic or contemporary EV-D68 isolates requires inoculation into either neonates or immunocompromised adults (Hixon, Yu, et al., 2017; Hurst et al., 2019; Morrey et al., 2018; Zhang, Zhang, Dai, et al., 2018). With each of these mouse models, the incidence and onset of AFM following infection is based on EV-D68 isolate and route of infection. Studies using these mouse models have been instrumental in demonstrating the increased virulence of contemporary versus historic EV-D68 isolates, and have served as platforms for demonstrating efficacy of candidate vaccines and antiviral treatments (Evans et al., 2019; Hixon, Clarke, et al., 2017; Vogt et al., 2020; Zhang, Zhang, Zhang, et al., 2018; Zheng et al., 2018). In a single published report, ferrets were shown to be permissive to the prototype EV-D68 Fermon strain, but respiratory infection did not result in significant respiratory illness, nor any evidence of AFM symptoms (Zheng et al., 2017). Similar to EV-D68 infection in ferrets, infection of cotton rats with either EV-D68 Fermon, EV-D68 VANBT/1 (pre-outbreak isolate) or EV-D68 MO/14/19 (outbreak isolate) revealed productive respiratory infection without any signs of AFM symptoms. Moreover, viral replication in the cotton rat is reported to be very rapid, with peak viral titers detected at 10 hours post-infection and viral clearance by 48 hours post-infection (Patel et al., 2016).

Nonhuman primates (NHPs) are the most phylogenetically similar preclinical species to humans, and as such, represent the gold standard for modelling human physiology and pathology, and for predicting human pharmacologic and toxicologic responses to candidate therapeutic compounds. The relatively large size of NHPs as compared with rodent models provides the additional advantage of increased sample collection volumes and frequencies, as well as the feasibility of collecting additional types of samples that mirror clinical sampling protocols. For example, viral challenge studies in nonhuman primates permit serial sampling of various biological compartments (e.g., bronchoalveolar lavage fluid) to characterize viral replication kinetics in sample types with direct clinical relevance (Vermillion et al., 2021). Moreover, advanced imaging techniques (e.g., MRI) and electromyography protocols are established for NHPs, providing additional translational utility for localizing spinal cord lesions and characterizing the patterns of denervation associated with AFM that can be correlated with human clinical cases.

Relevant to EV-D68, NHPs have been shown to harbor many different simian enteroviruses, with up to 72% amino acid identity to related human enteroviruses (Nix et al., 2008). Further, NHPs have been instrumental in the study of poliovirus infection and associated disease, as both rhesus and cynomolgus macaques have been shown to be susceptible to poliovirus infection and associated poliomyelitis following oral viral inoculation (Shen et al., 2017). Experimental infection of NHPs with EV-D68 is limited to a study in rhesus macaques, in which intranasal infection with EV-D68 (Kunming Strain) resulted in detectable virus in serially collected nasal swabs, blood, and fecal specimens out to 14 days post-infection (Zheng et al., 2018). Rhesus macaques have also been shown to mount potent neutralizing responses following immunization with an inactivated EV-D68 vaccine (Zheng et al., 2020). The susceptibility of other nonhuman primate species to EV-D68 infection and comparisons of nonhuman primate susceptibility to contemporary EV-D68 isolates, however, has not yet been evaluated.

In this study, juvenile NHPs – cynomolgus macaques, rhesus macaques, pigtailed macaques, and African green monkeys – were inoculated intratracheally and intranasally with one of five different 2014 or 2018 EV-D68 isolates (US/MO/2014-18947, US/IL/2014-18952, US/MD/2018-23209, US/MN/2018-23263, and US/WA/2018-23201) (**Table 1**). We report the results of clinical respiratory and neurologic monitoring, viral detection in nasal swabs, bronchoalveolar lavage and cerebrospinal fluid, and histopathology of respiratory and central nervous system tissues (**Fig. 1**). These collective data provide the first characterization of EV-D68 infection in NHP model systems.

**Table 1.**
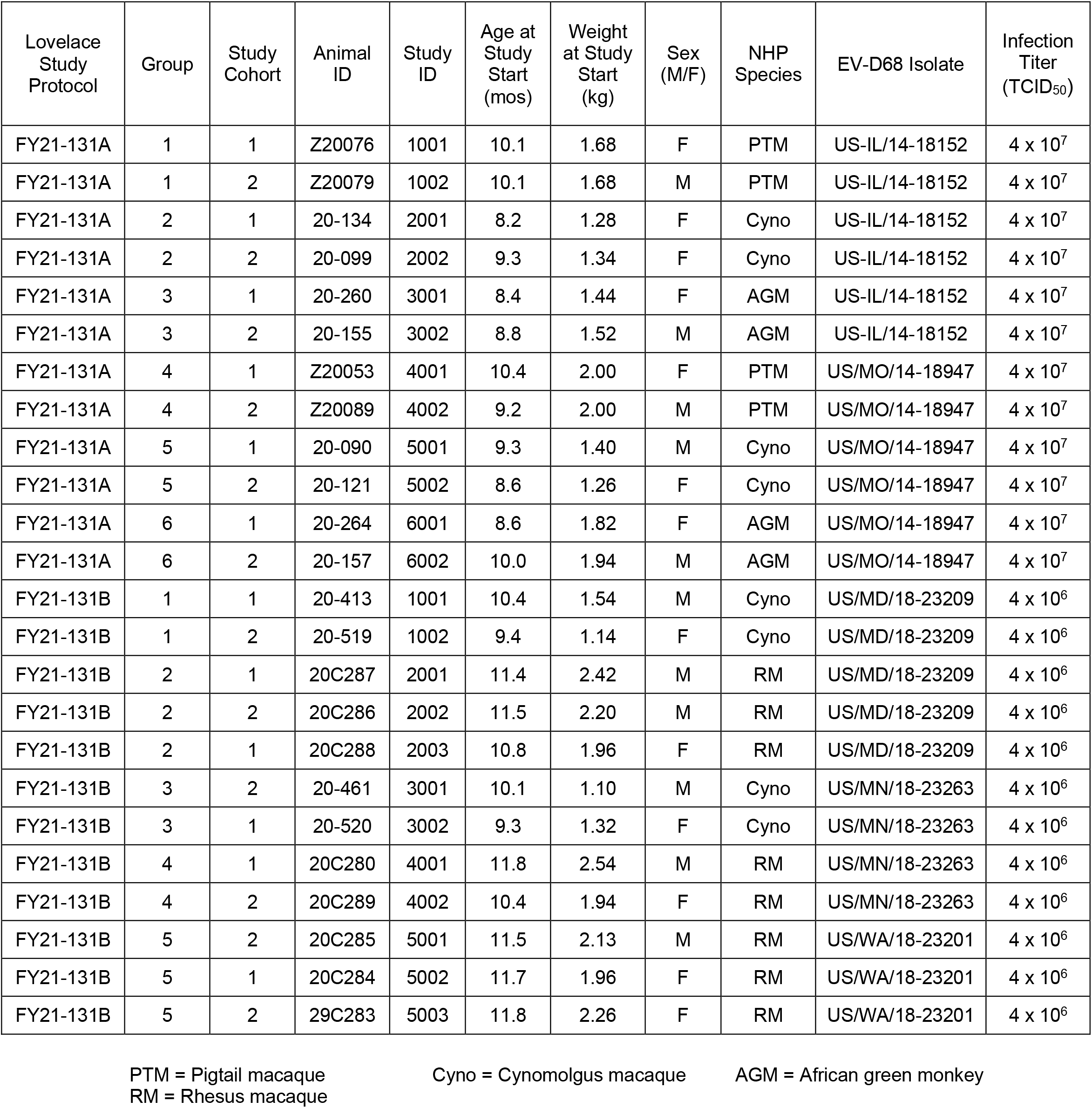
Animal Information and Group Assignments

**Figure 1.**
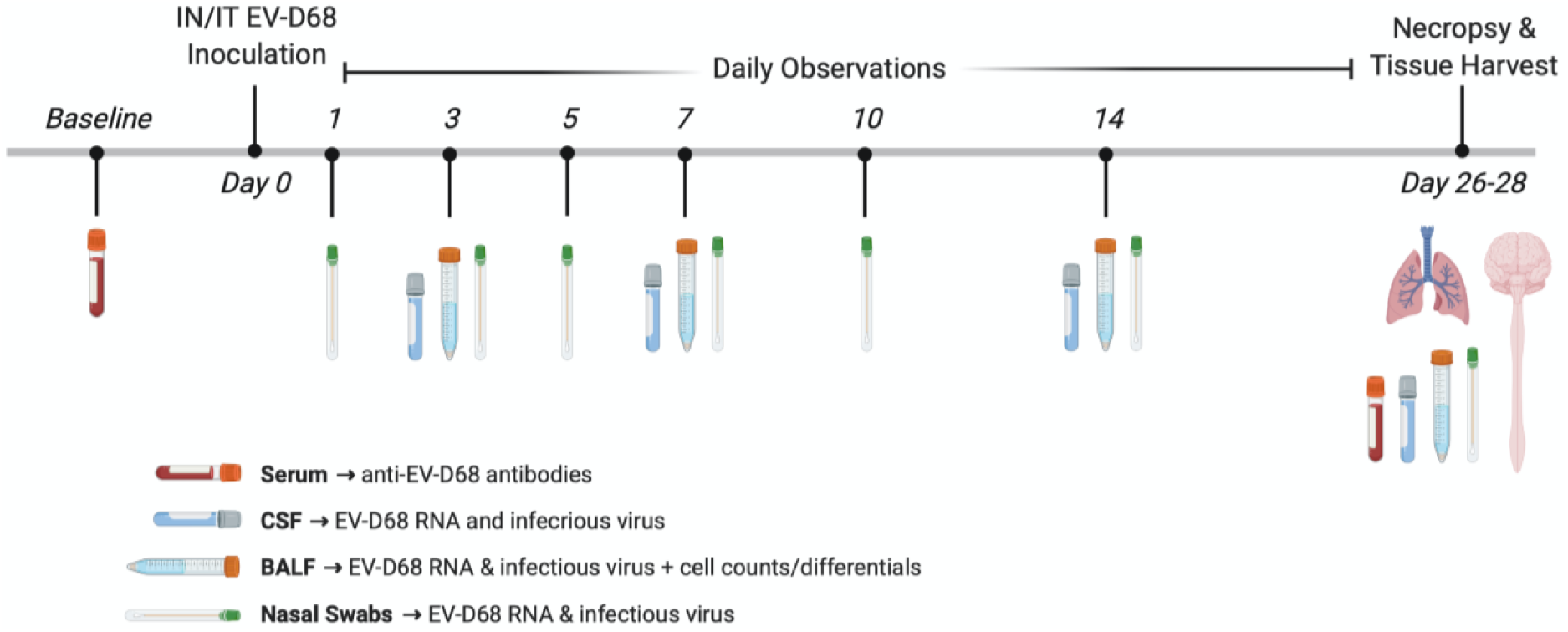
Experimental design and timeline. On study day 0, juvenile NHPs were inoculated with one of five different EV-D68 isolates by the respiratory route (n=2-3 per NHP species per EV-D68 isolate). Animals were monitored daily for evidence of clinical disease. Nasal swabs were collected 1, 3, 5, 7, 10, 14 and 26-28 days post-infection (dpi). Bronchoalveolar lavage fluid (BALF) and cerebrospinal fluid (CSF) were collected 3, 7, 14 and 26-28 dpi. EV-D68 RNA and infectious virus was quantified from nasal swabs, BALF and CSF by reverse-transcriptase quantitative PCR (RT-qPCR) and median tissue culture infectious dose (TCID_50_) assay, respectively. Cell counts and differentials were quantified from BALF cell pellets. Anti-EV-D68 antibodies were measured in serum collected at baseline and 26 dpi. Terminally collected respiratory tract and central nervous system tissues were formalin fixed and evaluated by histopathology.

## Results

### EV-D68 infection resulted in mild and sporadic clinical disease in all NHP species

Following EV-D68 infection, animals were monitored daily for clinical respiratory, gastrointestinal and neurological abnormalities. Body weights were obtained concurrent with anesthetized sample collection procedures. For all animals, regardless of the EV-D68 strain, clinical observations were limited to mild gastrointestinal and respiratory signs, and no neurologic abnormalities were observed from any animal during the observation period in this study (**Table 2**). For animals infected with the 2014 EV-D68 isolates, the majority of clinical observations were made 8 days post-infection (dpi) or later, and abnormal feces (i.e., soft stool or diarrhea) were the most commonly reported clinical sign (**Fig. 2a**), and nasal and/or ocular discharge was reported sporadically (**Fig. 2b**). Compared with cynomolgus macaques and pigtailed macaques, African green monkeys infected with either 2014 EV-D68 isolate had fewer clinical signs reported. There were no obvious differences in disease incidence or character when comparing infection with the IL/2014 and MO/2014 EV-D68 isolates in any NHP species (**Fig. 2a** and **b**). For animals infected with the 2018 EV-D68 isolates, respiratory symptoms were absent over the observation period for this study (**Fig. 2b**). Gastrointestinal symptoms were reported commonly in cynomolgus macaques, but these observations were also made at baseline, which suggests that they were unrelated to the EV-D68 infection (**Fig. 2a**). For both the 2014 and 2018 EV-D68 isolates, evidence of paresis, paralysis, or other neurologic disease was absent in all animals.

**Table 2.**
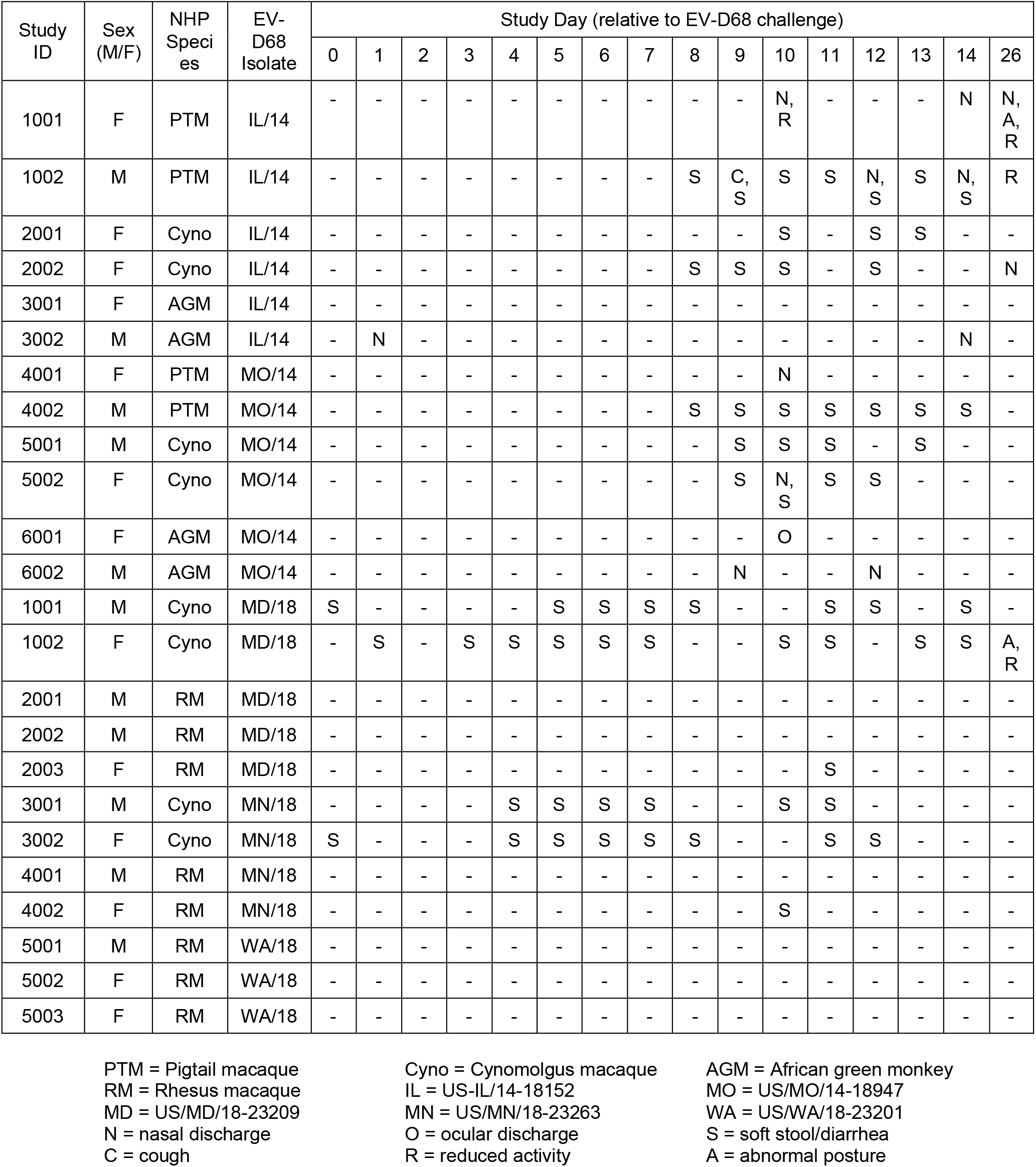
Clinical Observations

| Study ID | Sex (M/F) | NHP Species | EV-D68 Isolate | Study Day (relative to EV-D68 challenge) |  |  |  |  |  |  |  |  |  |  |  |  |  |  |  |
| --- | --- | --- | --- | --- | --- | --- | --- | --- | --- | --- | --- | --- | --- | --- | --- | --- | --- | --- | --- |
|  |  |  |  | 0 | 1 | 2 | 3 | 4 | 5 | 6 | 7 | 8 | 9 | 10 | 11 | 12 | 13 | 14 | 26 |
| 1001 | F | PTM | IL/14 | - | - | - | - | - | - | - | - | - | - | N, R | - | - | - | N | N, A, R |
| 1002 | M | PTM | IL/14 | - | - | - | - | - | - | - | - | S | C, S | S | S | N, S | S | N, S | R |
| 2001 | F | Cyno | IL/14 | - | - | - | - | - | - | - | - | - | - | S | - | S | S | - | - |
| 2002 | F | Cyno | IL/14 | - | - | - | - | - | - | - | - | S | S | S | - | S | - | - | N |
| 3001 | F | AGM | IL/14 | - | - | - | - | - | - | - | - | - | - | - | - | - | - | - | - |
| 3002 | M | AGM | IL/14 | - | N | - | - | - | - | - | - | - | - | - | - | - | - | N | - |
| 4001 | F | PTM | MO/14 | - | - | - | - | - | - | - | - | - | - | N | - | - | - | - | - |
| 4002 | M | PTM | MO/14 | - | - | - | - | - | - | - | - | S | S | S | S | S | S | S | - |
| 5001 | M | Cyno | MO/14 | - | - | - | - | - | - | - | - | - | S | S | S | - | S | - | - |
| 5002 | F | Cyno | MO/14 | - | - | - | - | - | - | - | - | - | S | N, S | S | S | - | - | - |
| 6001 | F | AGM | MO/14 | - | - | - | - | - | - | - | - | - | - | O | - | - | - | - | - |
| 6002 | M | AGM | MO/14 | - | - | - | - | - | - | - | - | - | N | - | - | N | - | - | - |
| 1001 | M | Cyno | MD/18 | S | - | - | - | - | S | S | S | S | - | - | S | S | - | S | - |
| 1002 | F | Cyno | MD/18 | - | S | - | S | S | S | S | S | - | - | S | S | - | S | S | A, R |
| 2001 | M | RM | MD/18 | - | - | - | - | - | - | - | - | - | - | - | - | - | - | - | - |
| 2002 | M | RM | MD/18 | - | - | - | - | - | - | - | - | - | - | - | - | - | - | - | - |
| 2003 | F | RM | MD/18 | - | - | - | - | - | - | - | - | - | - | - | S | - | - | - | - |
| 3001 | M | Cyno | MN/18 | - | - | - | - | S | S | S | S | - | - | S | S | - | - | - | - |
| 3002 | F | Cyno | MN/18 | S | - | - | - | S | S | S | S | S | - | - | S | S | - | - | - |
| 4001 | M | RM | MN/18 | - | - | - | - | - | - | - | - | - | - | - | - | - | - | - | - |
| 4002 | F | RM | MN/18 | - | - | - | - | - | - | - | - | - | - | S | - | - | - | - | - |
| 5001 | M | RM | WA/18 | - | - | - | - | - | - | - | - | - | - | - | - | - | - | - | - |
| 5002 | F | RM | WA/18 | - | - | - | - | - | - | - | - | - | - | - | - | - | - | - | - |
| 5003 | F | RM | WA/18 | - | - | - | - | - | - | - | - | - | - | - | - | - | - | - | - |

PTM = Pigtail macaque RM = Rhesus macaque MD = US/MD/18-23209 N = nasal discharge C = cough
Cyno = Cynomolgus macaque IL = US-IL/14-18152 MN = US/MN/18-23263 O = ocular discharge R = reduced activity
AGM = African green monkey MO = US/MO/14-18947 WA = US/WA/18-23201 S = soft stool/diarrhea A = abnormal posture

**Figure 2.**
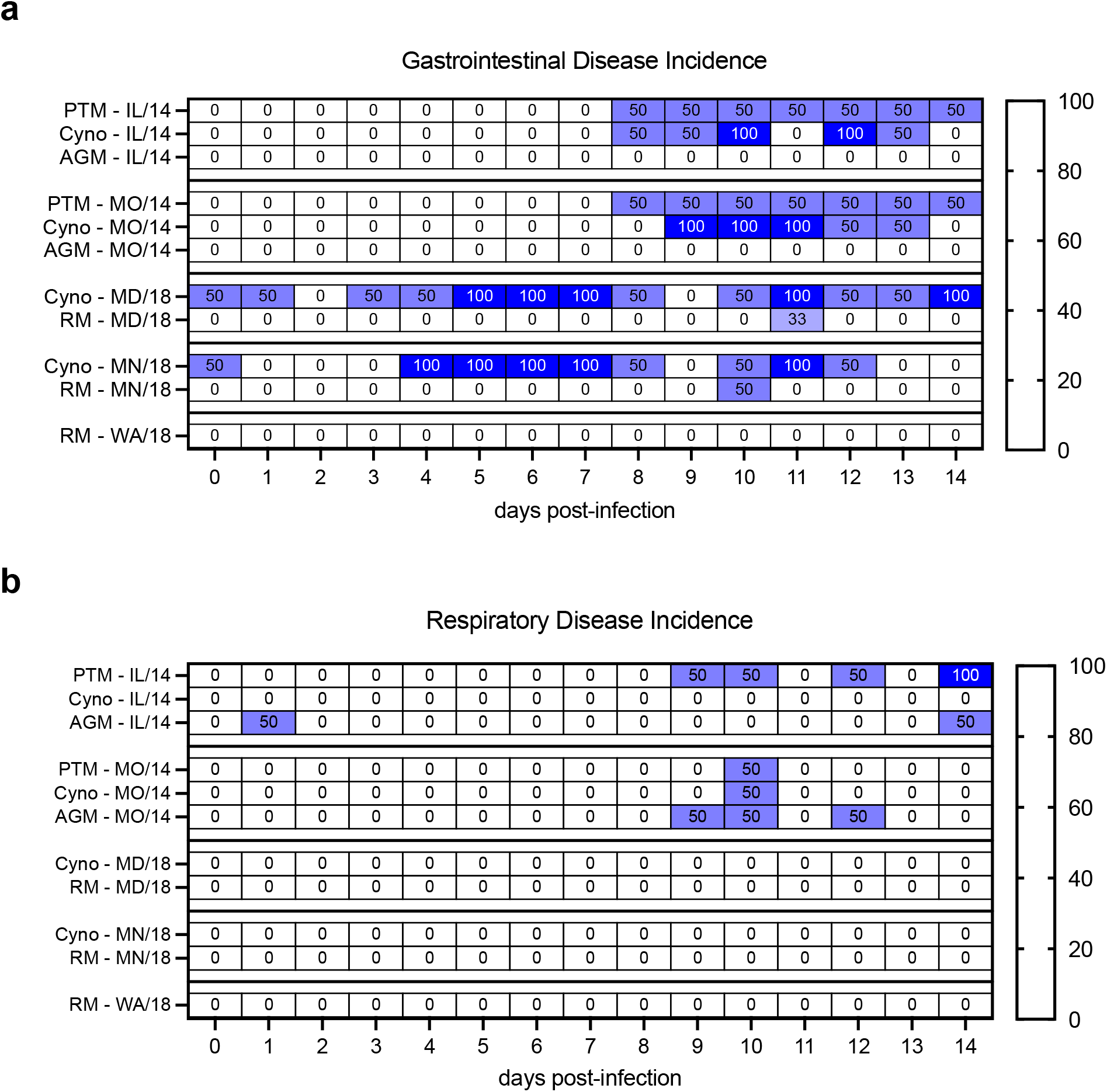
Clinical disease following EV-D68 infection of juvenile NHPs was mild and sporadic. Juvenile NHPs were inoculated with one of five different EV-D68 isolates by the respiratory route (n=2-3 per NHP species per EV-D68 isolate). Animals were monitored daily for evidence of clinical disease, and observations were categorized as either **a)** gastrointestinal (e.g., diarrhea or soft stool) or **b)** respiratory (e.g., nasal or ocular discharge). There were no observations associated with acute flaccid myelitis or other neurologic disease (e.g., paresis or paralysis). The incidence of each observation within each experimental group was calculated as a percentage for each observation day. PTM = pigtailed macaque; Cyno = cynomolgus macaque; AGM = African green monkey; RM = rhesus macaque.

For all infection groups, average change in body weight over the course of the study was less than 5% of baseline for most animals. Mild body weight loss was observed in some animals during the first week of the study, but was not consistently reported. Overall, animals gained weight over the course of the study, and the average increase in body weight was consistent with published growth curves for juvenile NHPs (Haertel et al., 2018). There was no obvious impact of either NHP species or EV-D68 isolate on body weight changes observed in this study (**Fig. 3a-d**). The absence of uninfected control animals in this study, however, precludes differentiation of any body weight loss or gain that may have been associated with collection procedures or normal expected growth, respectively.

**Figure 3.**
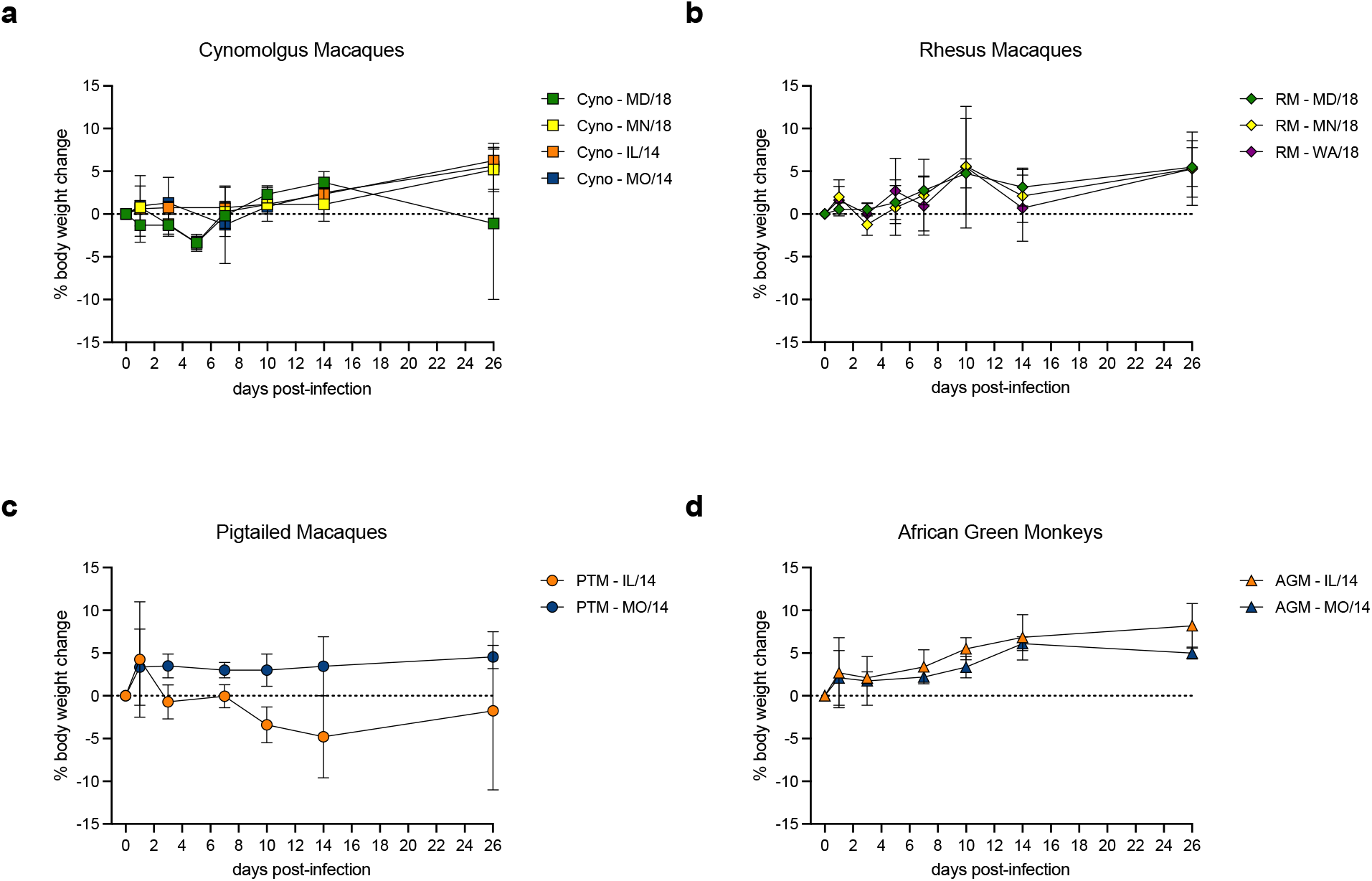
EV-D68 infection of juvenile NHPs resulted in minimal or no weight loss. Juvenile cynomolgus macaques (**a**), rhesus macaques (**b**), pigtailed macaques (**c**), and African green monkeys (**d**) were inoculated with one of five different EV-D68 isolates by the respiratory route (n=2-3 per NHP species per EV-D68 isolate). Body weights were collected concurrent with anesthetized procedures up to 26 days post-infection. The percent body weight change from baseline was calculated for each animal, and the mean +/- the standard error are shown for each group. PTM = pigtailed macaque; Cyno = cynomolgus macaque; AGM = African green monkey; RM = rhesus macaque.

### EV-D68 infection resulted in limited and transient detection of viral RNA in the upper and lower respiratory tract

In order to assess viral load and viral replication kinetics in the respiratory tract and central nervous system, bronchoalveolar lavage fluid (BALF), nasal swabs, and cerebrospinal fluid (CSF) were serially collected following EV-D68 inoculation (**Fig. 1**). Viral RNA was measured by RT-PCR and infectious virus was detected by median tissue culture infectious dose (TCID_50_) assay. In animals infected with the 2014 EV-D68 isolates, neither viral RNA nor infectious virus could be quantified in any of the samples analyzed (**Table 3**). In animals infected with the 2018 EV-D68 isolates, viral RNA was variably detected within nasal swabs and BALF supernatant collected between 1-5 dpi (**Fig. 4**). In nasal swabs collected 1 dpi, average EV-D68 RNA was quantified between 10^4^-10^6^ copies per swab for each virus isolate, and RNA load in nasal swabs was similar between rhesus (**Fig. 4a**) and cynomolgus (**Fig. 4c**) macaques. By 3 dpi, viral RNA was below the limit of quantification in nasal swabs collected from all animals except one rhesus macaque challenged with EV-D68 MD/2018 (**Fig. 4a** and **c**). In BALF collected 3 dpi, viral RNA was detectable in one rhesus from each the MD/2018 and WA/2018 infection groups (**Fig. 4b**), and from one cynomolgus macaque from the MN/2018 infection group (**Fig. 4d**). All BALF samples collected 7 dpi were below the assay limit of quantification (**Fig. 4b** and **d**). EV-D68 RNA was not detectable in CSF collected 3 dpi (**Table 3**). All samples evaluated by PCR for viral RNA were also tested for the presence of infectious virus but no infectious virus was detected (data not shown).

**Table 3.** EV-D68 RNA Concentrations and Western Blot Results

| Animal ID | Sex (M/F) | NHP Species | EV-D68 Isolate | EV-D68 RNA (copies/swab or copies/mL) |  |  |  |  |  |  | Western Blot Results |
| --- | --- | --- | --- | --- | --- | --- | --- | --- | --- | --- | --- |
|  |  |  |  | Nasal Swabs |  |  |  | BALF |  | CSF |  |
|  |  |  |  | 1 dpi | 3 dpi | 5 dpi | 7 dpi | 3 dpi | 7 dpi | 3 dpi |  |
| 1001 | F | PTM | IL | BQL | BQL | BQL | BQL | BQL | BQL | BQL | NEG |
| 1002 | M | PTM | IL | BQL | BQL | BQL | BQL | BQL | BQL | BQL | NEG |
| 2001 | F | Cyno | IL | BQL | BQL | BQL | BQL | BQL | BQL | BQL | POS |
| 2002 | F | Cyno | IL | BQL | BQL | BQL | BQL | BQL | BQL | BQL | NEG |
| 3001 | F | AGM | IL | BQL | BQL | BQL | BQL | BQL | BQL | BQL | NEG |
| 3002 | M | AGM | IL | BQL | BQL | BQL | BQL | BQL | BQL | BQL | NEG |
| 4001 | F | PTM | MO | BQL | BQL | BQL | BQL | BQL | BQL | BQL | POS |
| 4002 | M | PTM | MO | BQL | BQL | BQL | BQL | BQL | BQL | BQL | NEG |
| 5001 | M | Cyno | MO | BQL | BQL | BQL | BQL | BQL | BQL | BQL | POS |
| 5002 | F | Cyno | MO | BQL | BQL | BQL | BQL | BQL | BQL | BQL | NEG |
| 6001 | F | AGM | MO | BQL | BQL | BQL | BQL | BQL | BQL | BQL | NEG |
| 6002 | M | AGM | MO | BQL | BQL | BQL | BQL | BQL | BQL | BQL | NEG |
| 1001 | M | Cyno | MD | 8.40E+04 | BQL | BQL | BQL | BQL | BQL | BQL | POS |
| 1002 | F | Cyno | MD | 2.35E+03 | BQL | BQL | BQL | BQL | BQL | BQL | POS |
| 2001 | M | RM | MD | 8.71E+04 | BQL | BQL | BQL | BQL | BQL | BQL | NEG |
| 2002 | M | RM | MD | 1.89E+05 | 1.92E+04 | 1.17E+03 | BQL | BQL | BQL | BQL | NEG |
| 2003 | F | RM | MD | 8.58E+02 | BQL | BQL | BQL | 1.77E+04 | BQL | BQL | POS |
| 3001 | M | Cyno | MN | 4.72E+03 | BQL | BQL | BQL | 6.58E+03 | BQL | BQL | POS |
| 3002 | F | Cyno | MN | BQL | BQL | BQL | BQL | BQL | BQL | BQL | POS |
| 4001 | M | RM | MN | 1.07E+03 | BQL | BQL | BQL | BQL | BQL | BQL | POS |
| 4002 | F | RM | MN | 1.57E+04 | BQL | BQL | BQL | BQL | BQL | BQL | NEG |
| 5001 | M | RM | WA | 9.23E+04 | BQL | BQL | BQL | 6.05E+03 | BQL | BQL | POS |
| 5002 | F | RM | WA | BQL | BQL | BQL | BQL | BQL | BQL | BQL | NEG |
| 5003 | F | RM | WA | 2.55E+04 | BQL | BQL | BQL | BQL | BQL | BQL | NEG |

PTM = Pigtail macaque RM = Rhesus macaque MD = US/MD/18-23209 DPI = days post-infection
Cyno = Cynomolgus macaque IL = US-IL/14-18152 MN = US/MN/18-23263 BQL = below the quantification limit of the assay
AGM = African green monkey MO = US/MO/14-18947 WA = US/WA/18-23201

**Figure 4.**
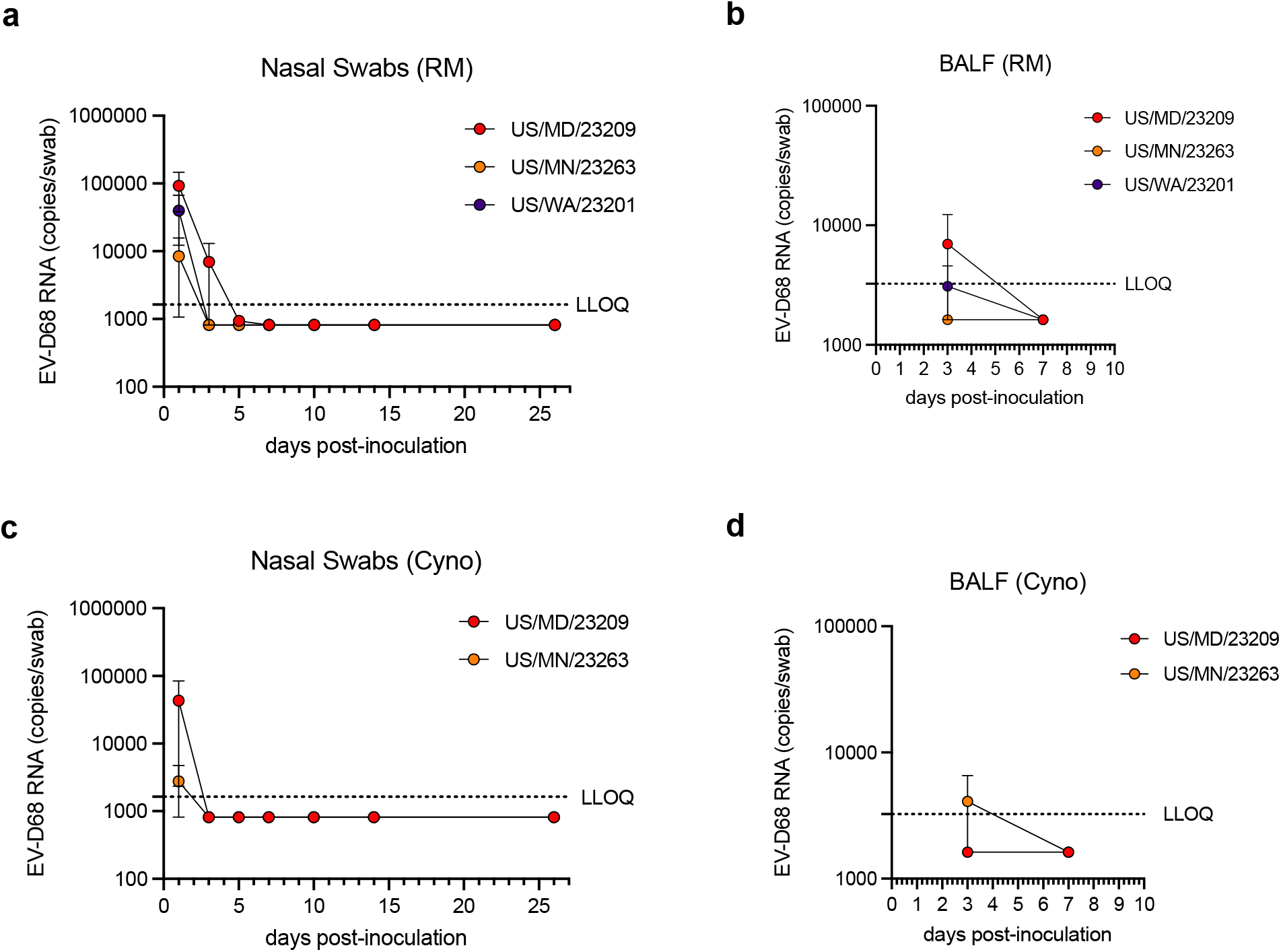
EV-D68 2018 infection resulted in limited and transient detection of viral RNA in the upper and lower respiratory tract. Juvenile rhesus and cynomolgus macaques were inoculated with one of three different 2018 EV-D68 isolates by the respiratory route (n=2-3 per NHP species per EV-D68 isolate). Nasal swabs (**a** and **c**) and bronchoalveolar lavage fluid (BALF, **b** and **d**) were serially collected from each animal up to 26 days post-infection (dpi), and EV-D68 RNA was quantified by RT-qPCR. RNA copies per swab or copies per mL were calculated for each sample, and the mean +/- the standard error are shown for each group. Cyno = cynomolgus macaque; RM = rhesus macaque. LLOQ = lower limit of quantification.

Because EV-D68 RNA was only transiently detected in upper and lower respiratory tract samples, western blot for anti-EV-D68 antibodies was performed on serum collected at baseline and at 26 dpi in order to confirm infection (**Fig. 5a**). Other than animal #20-413 (cynomolgus macaque from the MD/2018 infection group), for which baseline antibody screening was inconclusive (data not shown), all baseline serum samples were negative for anti-EV-D68 antibodies, confirming no pre-existing immunity to EV-D68 (**Table 3**). Overall, infection with the 2018 EV-D68 isolates resulted in the greatest seroconversion rate, with 100% of the cynomolgus macaques and between 33-50% of the rhesus macaques having detectable EV-D68 antibodies. Seroconversion for the 2014 EV-D68 isolates was overall lower, ranging from 0-50%. Notably, there was no evidence of seroconversion in African green monkeys infected with either 2014 EV-D68 isolate (**Fig. 5b**).

**Figure 5.**
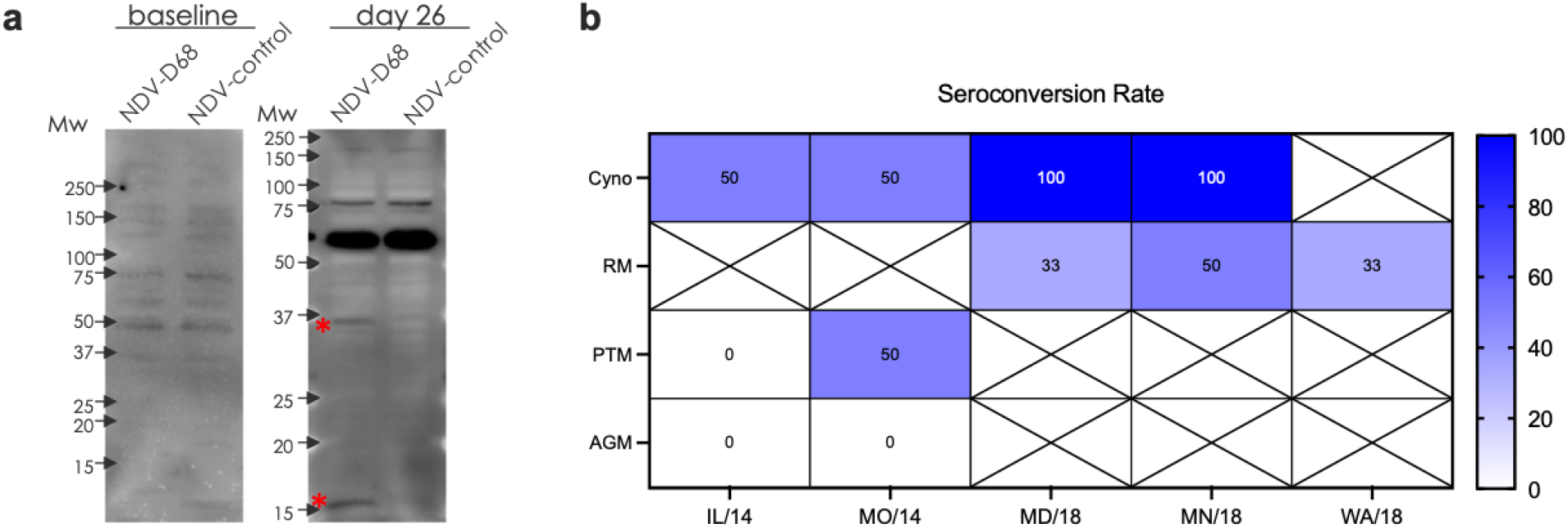
Anti-EV-D68 antibodies were detectable in the serum of juvenile NHPs following respiratory challenge with contemporary EV-D68 isolates. Juvenile cynomolgus macaques, rhesus macaques, pigtailed macaques, and African green monkeys were inoculated with one of five different EV-D68 isolates by the respiratory route (n=2-3 per NHP species per EV-D68 isolate). Blood was collected at baseline and at 26 days post-infection (dpi) for detection of serum antibodies against the EV-D68 structural proteins by western blot assay. A representative western blot shows positive bands at ∼35 and ∼15 kDa (asterisks), which represent antibodies against EV-D68 capsid proteins at 26 dpi (**a**). The seroconversion rate was calculated as a percentage for each experimental group (**b**). PTM = pigtailed macaque; Cyno = cynomolgus macaque; AGM = African green monkey; RM = rhesus macaque.

Interestingly, the presence of anti-EV-D68 antibodies did not always correlate with the presence of detectable EV-D68 RNA in respiratory samples; there were 4 animals (3 cynomolgus macaques and 1 pigtailed macaque) with detectable antibodies at 26 dpi that had no detectable virus in either nasal swabs or BALF (**Table 3**). This suggests that there was productive EV-D68 infection that was either not captured within the sampling window or was below the limit of detection of the PCR assay. Surprisingly, there were also 4 animals (all rhesus macaques) with detectable EV-D68 RNA in nasal swabs collected 1 dpi, but no evidence of serum antibodies at 26 dpi (**Table 3**). It is possible that viral RNA detected in the nasal cavity 1 dpi may represent residual RNA from the EV-D68 inoculum; animal #2002, however, also had detectable viral RNA in nasal swabs collected 3 and 5 dpi, which makes this hypothesis less likely. This disparity may also be explained by species-specific differences in immune response to EV-D68 infection.

### EV-D68 infection resulted in mild airway inflammation without evidence of virus-induced pulmonary histopathology

In order to characterize the airway inflammation following EV-D68 infection, BALF cell pellets were processed and analyzed for total cell counts and cell differentials at each collection timepoint (**Fig. 1**). Due to species-specific variability in the baseline total cell counts in BALF, data were normalized and presented as a percent change from baseline for each post-infection timepoint (**Fig. 6**). Compared with macaques (**Fig. 6a-c**), AGM demonstrated greater overall increases in airway inflammation post-infection, regardless of EV-D68 isolate, though response was highly variable (**Fig. 6d**). This inflammatory response was observed independent of any evidence of EV-D68 infection, based on absence of serum antibodies, suggesting that this response may represent non-specific inflammation related to the BALF collection procedures. Differentiation of procedural-related changes cannot be made without an uninfected control group for comparison. Comparing infection of different EV-D68 isolates within NHP species, EV-D68 MD/2018 resulted in a greater induction of airway inflammation than the other EV-D68 isolates evaluated in both cynomolgus macaques and rhesus macaques at 26 dpi (**Fig. 6a** and **b**). Overall, pigtailed macaques showed the smallest increase in airway inflammatory cells over the course of the study (**Fig. 6c**).

**Figure 6.**
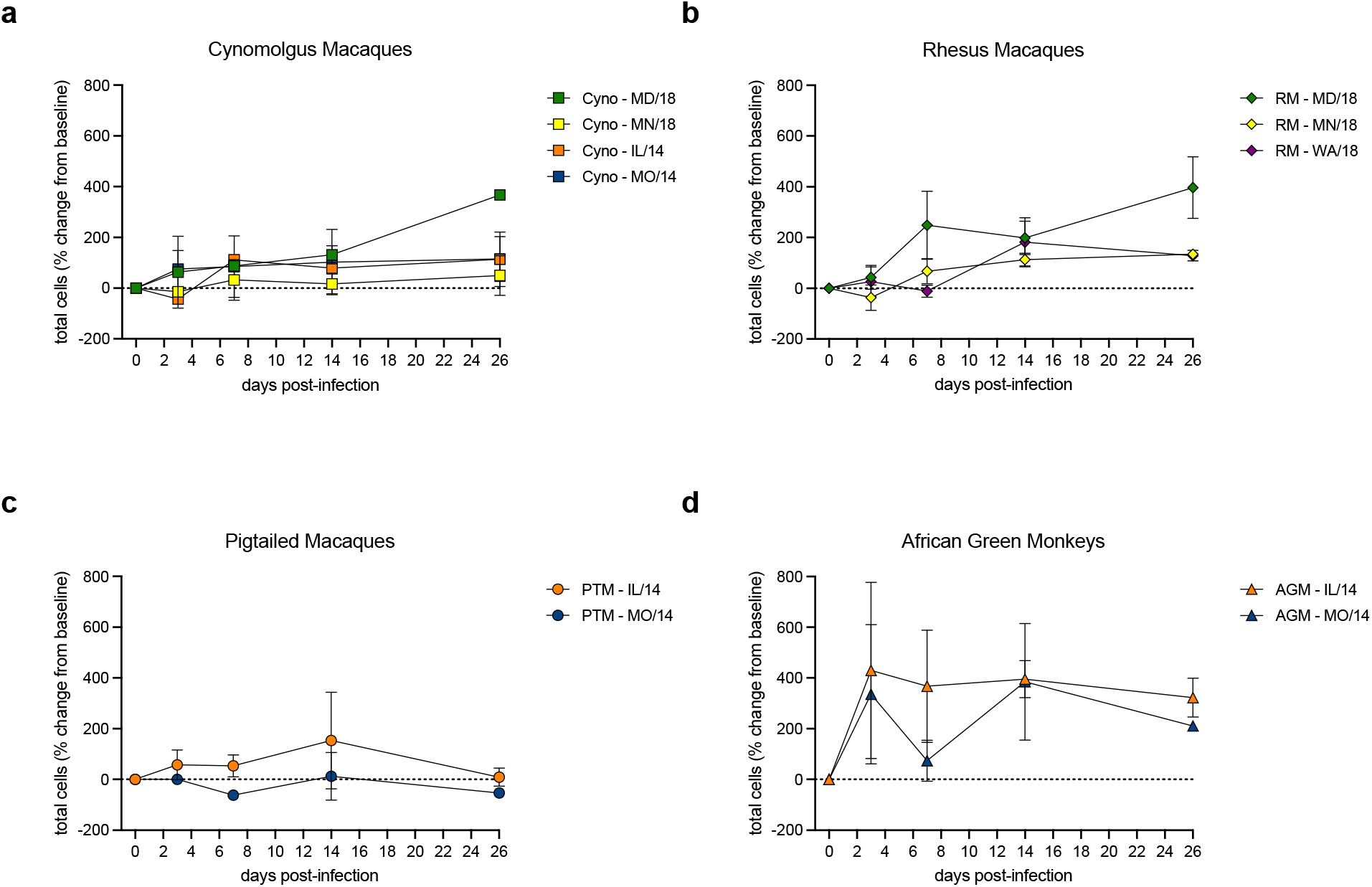
Airway inflammation following EV-D68 infection of juvenile NHPs was species and strain specific. Juvenile cynomolgus macaques (**a**), rhesus macaques (**b**), pigtailed macaques (**c**), and African green monkeys (**d**) were inoculated with one of five different EV-D68 isolates by the respiratory route (n=2-3 per NHP species per EV-D68 isolate). Bronchoalveolar lavage fluid (BALF) was serially collected from each animal up to 26 days post-infection (dpi), and total live cells were quantified from BALF cytospins. The percent change from baseline in total cells was calculated for each sample, and the mean +/- the standard error are shown for each group. PTM = pigtailed macaque; Cyno = cynomolgus macaque; AGM = African green monkey; RM = rhesus macaque.

At 26-28 dpi, animals were euthanized for terminal tissue collections. Terminal lung weights were obtained and normalized to terminal body weights and brain weight, and ratios were calculated for each animal (**Table 4**). Overall, terminal lung weights ranged from 0.6-0.9% of total body weight, and from 12-22% of total brain weight. Some species-specific differences in lung/body weight and lung/brain weight ratios were observed, but within NHP species, there were no clear differences in lung weight ratios between EV-D68 isolates. The absence of age-matched uninfected controls for each NHP species precludes comparison of post-infection organ weight ratios with normal ranges, though the reported lung to body weight ratios are comparable to published reference intervals for NHPs (Miller et al., 2017), suggesting an absence of any significant pulmonary inflammation or edema in lungs collected 26-28 dpi. Sections of the thoracic trachea, left lungs, tracheobronchial lymph nodes, brain and spinal cord were microscopically examined and assessed for evidence of inflammation and/or injury. No evidence of virus-associated pathology was noted for any of the tissues evaluated in this study (data not shown).

**Table 4.** Terminal Lung Weight Ratios

| Animal ID | Sex (M/F) | NHP Species | EV-D68 Isolate | Terminal Body Weight (kg) | Terminal Lung Weight (g) | Terminal Brain Weight (g) | Terminal Lung/Body Weight Ratio (g/kg) | Terminal Lung/Brain Weight Ratio (g/g) |
| --- | --- | --- | --- | --- | --- | --- | --- | --- |
| 1001 | F | PTM | IL | 1.72 | 15.43 | 92.06 | 8.973 | 0.168 |
| 1002 | M | PTM | IL | 1.74 | 11.71 | 81.34 | 6.729 | 0.144 |
| 2001 | F | Cyno | IL | 1.28 | 9.94 | 61.14 | 7.762 | 0.163 |
| 2002 | F | Cyno | IL | 1.44 | 10.62 | 62.15 | 7.376 | 0.171 |
| 3001 | F | AGM | IL | 1.52 | 11.24 | 61.59 | 7.393 | 0.182 |
| 3002 | M | AGM | IL | 1.64 | 11.99 | 62.66 | 7.309 | 0.191 |
| 4001 | F | PTM | MO | 1.94 | 12.12 | 79.45 | 6.245 | 0.152 |
| 4002 | M | PTM | MO | 2.06 | 14.69 | 90.35 | 7.132 | 0.163 |
| 5001 | M | Cyno | MO | 1.42 | 9.97 | 73.10 | 7.018 | 0.136 |
| 5002 | F | Cyno | MO | 1.30 | 9.55 | 62.12 | 7.347 | 0.154 |
| 6001 | F | AGM | MO | 1.84 | 12.42 | 60.60 | 6.751 | 0.205 |
| 6002 | M | AGM | MO | 1.96 | 13.04 | 65.94 | 6.652 | 0.198 |
| 1001 | M | Cyno | MD | 1.66 | 13.88 | 66.94 | 8.364 | 0.207 |
| 1002 | F | Cyno | MD | 1.08 | 7.70 | 54.60 | 7.132 | 0.141 |
| 2001 | M | RM | MD | 2.50 | 17.79 | 89.41 | 7.114 | 0.199 |
| 2002 | M | RM | MD | 2.28 | 16.23 | 88.42 | 7.119 | 0.184 |
| 2003 | F | RM | MD | 2.14 | 13.85 | 70.38 | 6.471 | 0.197 |
| 3001 | M | Cyno | MN | 1.18 | 7.39 | 58.32 | 6.263 | 0.127 |
| 3002 | F | Cyno | MN | 1.38 | 9.10 | 54.00 | 6.596 | 0.169 |
| 4001 | M | RM | MN | 2.74 | 16.70 | 89.73 | 6.095 | 0.186 |
| 4002 | F | RM | MN | 2.02 | 12.97 | 64.41 | 6.419 | 0.201 |
| 5001 | M | RM | WA | 2.26 | 15.17 | 84.12 | 6.714 | 0.180 |
| 5002 | F | RM | WA | 2.08 | 14.67 | 70.02 | 7.051 | 0.209 |
| 5003 | F | RM | WA | 2.38 | 16.00 | 75.25 | 6.723 | 0.213 |

PTM = Pigtail macaque RM = Rhesus macaque MD = US/MD/18-23209
Cyno = Cynomolgus macaque IL = US-IL/14-18152 MN = US/MN/18-23263
AGM = African green monkey MO = US/MO/14-18947 WA = US/WA/18-23201

## Discussion

While *in vitro* studies have significantly expanded our understanding of the mechanisms of EV-D68 cell entry and replication, confirmation of these concepts *in vivo* has yielded variable results (Hixon, Clarke, et al., 2017). Establishing and characterizing additional large animal models of EV-D68 infection and disease is critical for efficacy and safety evaluations for candidate medical countermeasures. Herein, we describe the results of pilot EV-D68 infection studies in juvenile NHPs, in which viral replication kinetics and clinical disease are characterized over 26-28 days following infection by the respiratory route.

In this study, inoculation with either 2014 or 2018 EV-D68 isolates by both the intranasal and intratracheal routes resulted in sporadic, mild and transient clinical disease that was characterized by gastrointestinal and/or upper respiratory symptoms that were most commonly observed 6-14 days post-infection. While the 2014 EV-D68 isolates resulted in sporadic respiratory signs (i.e., nasal or ocular discharge), infection with the 2018 EV-D68 isolates did not. No neurologic abnormalities were observed in any animal during the observation period in this study. Moreover, no gross or histopathological findings were noted in the respiratory tract or spinal cord that could be attributed to EV-D68 infection. Overall, clinical disease and pathology are not consistent components of this model, and are unlikely to serve as a useful markers of efficacy for evaluating candidate medical countermeasures. This outcome is consistent with observations from experimental infections of immunocompetent cotton rats and ferrets, in which clinical disease is minimal or absent (Patel et al., 2016; Zheng et al., 2017). Although the available surveillance data in humans is insufficient for determining the prevalence of clinically silent EV-D68 infections within the general population, it has been estimated that only 10% of EV-D68 respiratory infections resulting in hospitalization are at risk of developing AFM (Dyda et al., 2018; Morens et al., 2019). The absence of neurologic symptoms in this NHP infection study, therefore, is not an unexpected result based on the total number of animals enrolled.

Although the specific receptor(s) for EV-D68 remain unknown, sialic acid has been hypothesized, and studies *in vitro* have demonstrated that prototype EV-D68 and isolates from 2010-2011 bind both α2,6- and α2,3-linked sialic acids (Baggen et al., 2016; Imamura et al., 2014). EV-D68 strains isolated from 2012 and onward, however, have also demonstrated sialic acid-independent binding *in vitro* (Baggen et al., 2016; Liu et al., 2015), suggesting that more contemporary EV-D68 strains may use alternative cell receptors, which may influence tissue tropism and disease. One potential limitation of NHP models of respiratory infection is the differential expression and distribution of sialic acid receptors across different cell types within the upper and lower respiratory tract. Compared with humans, who express both α2,6- and α2,3-linked sialic acid residues in ciliated respiratory epithelial cells, sialic acid expression in NHPs is more limited, and receptors are present only on pulmonary Type II pneumocytes, submucosal goblet cells, and upper respiratory tract ciliated epithelium (Kuchipudi et al., 2021). Additional studies are needed to characterize receptor and co-receptor usage by both historic and contemporary EV-D68 isolates in order to predict how this impacts species-specific susceptibility to infection.

Indeed, much of the challenge with modeling the AFM phenotype secondary to a respiratory EV-D68 infection stems from the fact that the incidence of this specific disease sequelae is very low in humans. Likewise, even in the perfect model system, incidence of AFM is predicted to be low, which then obligates enrollment of very large cohorts of animals in order to detect the phenotype in a sufficient number to evaluate outcomes. The obvious workarounds to this dilemma are to artificially increase disease incidence through various manipulations of the host and/or virus. Such strategies have been employed in mouse models of EV-D68 infection, where productive viral replication and overt disease require either direct inoculation of the virus into the central nervous system of neonatal animals or the use of immunodeficient mouse strains. Future EV-D68 infection studies in NHPs should consider concurrent immunosuppression and/or direct intrathecal inoculation of virus in order to enhance infection and disease.

## Methods

### Study Design

A total of 24 animals were exposed to one of five EV-D68 clinical isolates by combined intranasal and intratracheal instillation. The study was conducted in two phases: Phase I (conducted under Lovelace protocol FY20-131A) evaluated the 2014 EV-D68 isolates in n=12 animals, and Phase II (conducted under Lovelace protocol FY20-131B) evaluated the 2018 EV-D68 isolates in n=12 animals. Within each study phase, animals were divided into two cohorts of n=6/cohort – staggered by one day – with even distribution of each study group across cohorts (**Table 1**). Animals were observed daily following inoculation, and blood, bronchoalveolar lavage fluid (BALF), nasal swabs, and cerebrospinal fluid (CSF) were serially collected for determination of viral replication kinetics (**Fig. 1**). Necropsy and terminal tissue collection was performed 26-28 days post-infection for histopathology.

### Viruses

The following EV-D68 isolates were evaluated in this study: US/MO/2014-18947 (Clade B1, GenBank Accession No. KM851225), US/IL/2014-18952 (Clade B2, GenBank Accession No. KM851230), US/MD/2018-23209 (Clade B3, GenBank Accession No. MN246002), US/MN/2018-23263 (Clade B3, GenBank Accession No. MN246026), and US/WA/2018-23201 (Clade B3, GenBank Accession No. MN245994). All virus stocks were sourced from ATCC, and all seed and working stocks for animal studies were generated by the Pekosz Laboratory at Johns Hopkins. Human rhabdomyosarcoma (RD) cells were obtained from the American Type Culture Collection (ATCC) (CCL-136) and cultured in Gibco Dulbecco’s Minimal Essential Medium (DMEM) supplemented with 10% fetal bovine serum (FBS), 100 U/mL penicillin, 100 μg/mL streptomycin, 1 mM sodium pyruvate and 2 mM GlutaMAX (Gibco). The cells were grown at 37°C in a humidified environment supplemented to 5% CO2. The initial vial of cells was expanded to obtain 20 vials of frozen cells which were confirmed to be mycoplasma free. One vial of this stock was thawed and expanded for use, with one additional stock of 20 frozen cells generated from the first passage after thawing. The cultures were passaged up to 20-25 times before new cells were thawed. Virus was stored at −70°C or below until infections. For IN/IT instillations, virus was thawed, diluted with Dulbecco’s Modified Eagle Medium (DMEM), and stored on ice until inoculations. Two aliquots of each inoculum were used for back-titration and confirmation of titer by TCID_50_ assay using RD cells. The inoculation stock titers for US/MO/2014-18947 and US/IL/2014-18952 were 2 × 10^7^ TCID_50_/mL, and the inoculation stock titers for US/MD/2018-23209, US/MN/2018-23263, and US/WA/2018-23201 were 2 × 10^6^ TCID_50_/mL. The VP gene of each virus stock was sequenced to confirm the isolate designation. All virus infections were performed at 32°C.

### Animals

All animal studies were conducted at Lovelace Biomedical Research Institute (LBRI, Albuquerque, NM). All infection studies were performed in an animal biosafety level 2 (ABSL-2) laboratory. All protocols were reviewed by an Institutional Animal Care and Use Committee (IACUC) at LBRI (IACUC #FY20-131). Research was conducted under an IACUC approved protocol in compliance with the Animal Welfare Act, PHS Policy, and federal statutes and regulations relating to animals and experiments involving animals. The facilities where this research was conducted are accredited by the Association for Assessment and Accreditation of Laboratory Animal Care, International and strictly adhere to principles stated in the Guide for the Care and Use of Laboratory Animals, National Research Council, 2011 (National Academies Press, Washington, DC). A total of 24 animals were used on this study [8 cynomolgus macaques (3M/5F); 8 rhesus macaques (4M/4F); 4 pigtailed macaques (2M/2F); and 4 African green monkeys (2M/2F)]. Cynomolgus macaques and African green monkeys were sourced from World Wide Primates (Miami, FL); rhesus macaques were sourced from Envigo (Alice, TX); and pigtailed macaques were sourced from the University of Washington National Primate Research Center (Mesa, AZ). Animals were between 8.2 and 11.8 months of age and between 1.1 and 2.5 kg at the time of EV-D68 inoculation (**Table 1**). All animals were quarantined for a minimum of 21 days following arrival at LBRI. All animals were tested and confirmed negative for *Mycobacterium tuberculosis*, simian immunodeficiency virus (SIV), simian T-lymphotropic virus-1 (STLV-1), simian retroviruses 1 and 2 (SRV-1 and SRV-2), and *Macacine herpesvirus* 1 (Herpes B virus). Prior to study start, all animals were examined by a laboratory animal veterinarian, and only healthy animals were placed on study. Animals were socially housed in a climate-controlled room with a fixed light/dark cycle (12-h light/12-h dark). Animals were monitored at least twice daily throughout the study. Commercial monkey chow, treats, and fruit were provided twice daily by trained personnel. Water was available ad libitum.

### Viral Inoculations

Animals were anesthetized with ketamine and xylazine and placed in the prone position for EV-D68 infections by both intranasal and intratracheal instillation. The total inoculation was delivered in 2 mL per animal (1 mL delivered intratracheal and 1 mL delivered intranasal – 0.5 mL per nostril). Virus was thawed on wet ice and diluted immediately prior to infections. For intratracheal instillation, a pediatric bronchoscope (Olympus XP-40) was advanced to the level of the carina. Polyethylene (PE) tubing was advanced through the bronchoscope, and 1 mL virus was instilled through the tubing, followed by 0.5 mL sterile saline and approximately 1 mL air. For intranasal instillation, the viral inoculum was administered drop-wise into each nostril using a calibrated pipette (i.e., 0.5 mL per nostril, 1 mL total).

### Clinical Scoring

Animals were observed at baseline and daily post-infection for evidence of clinical disease, including respiratory illness and paresis/paralysis. Respiratory signs were recorded based on a semi-quantitative scoring system comprised on observations of nasal or ocular discharge, cough, labored breathing, and abnormal posture. Signs of paresis or paralysis were recorded based on the number and severity of affected limbs, using an adapted scoring system (Urbanski et al., 2012). Severity of the paresis/paralysis was recorded on a scale of 0-3 (0 = normal motor function; 1 = mild paresis [e.g., slight favoring of contralateral limb and/or reduced gripping with hands/feet]; 2 = moderate paresis [e.g., obvious favoring of contralateral limb and/or reduced use of entire limb; 3 = complete limb paralysis) for each of the four limbs. Other detailed observations were recorded based on a standard lexicon. Body weights were also collected at baseline, day 0, and 1, 3, 5, 7, 10, 14, and 26 days post-infection.

### Blood Collection

Blood was collected at baseline and 26 days post-infection for detection of serum anti-EV-D68 antibodies by western blot. Blood was collected from the femoral, saphenous or cephalic veins, and was performed in sedated or anesthetized animals. Whole blood was collected into serum separator tubes (SST) at each scheduled timepoint. Blood collected into SST was centrifuged at 2500 x *g* for 10 minutes at room temperature for isolation of serum. Sample aliquots were stored at −70ºC or below until analysis.

### Nasal Swab Collection

Nasal swabs were collected for measurement of viral burden at baseline and 1, 3, 5, 7, 10, 14, and 26 days post-infection. Samples were collected from sedated or anesthetized animals. Nylon flocked swabs (FLOQSwab, COPAN Diagnostics, Inc.) were advanced into the nasal cavity, and both nostrils were sampled using a single swab. Two swabs were collected per site. One swab was placed in a tube containing approximately 0.5 mL sterile saline and immediately frozen on dry ice prior to storage at −70 to −90ºC. The second swab was placed in a tube containing 1 mL of Trizol, was allowed to set at room temperature for 15 minutes, and then frozen on dry ice prior to storage at −70 to −90ºC.

### Bronchoalveolar Lavage Fluid Collection

At baseline and 3, 7, 14, and 26 days post-infection, bronchoalveolar lavage fluid (BALF) was collected for quantification of viral load. BALF was collected using a pediatric bronchoscope (Olympus XP40) that was advanced approximately 3-4 generations into the airways of one side of the lung. Approximately 5 mL sterile saline was instilled and then aspirated. The site of sample collection alternated between the left and right lung at each sequential timepoint. The total recovered volume was recorded. Samples were stored on wet ice until processing, at which time they were centrifuged at approximately 1000 x *g* at 4–C for 10 minutes. The supernatants were aliquoted and stored at −70ºC or below until analysis. The cell pellets were resuspended in media, and total cell counts and cell differentials were performed on stained cytospins. The remaining dry cell pellet was stored at −70ºC.

### Cerebrospinal Fluid Collection

At baseline, 3, 7, 14, and 26 days post-infection, cerebrospinal fluid (CSF) was collected for quantification of viral load. Animals were anesthetized with ketamine and xylazine for collections. The back of the head was clipped and sterile prepped. Approximately 0.3 mL CSF was collected by sterile puncture of the cisterna magna using a 22 G, 1” needle. CSF was stored in two aliquots at −70ºC or below until analysis.

### Necropsy and Tissue Collection

At 26-28 days post-infection, animals were euthanized for terminal tissue collections. Terminal heart and lung weights were obtained and normalized to terminal body weights and brain weight, and ratios were calculated for each animal. Sections of the right lung were collected and flash frozen in liquid nitrogen for viral quantification assays (i.e., RT-qPCR and TCID_50_). All frozen samples were stored at −70 °C or below until processing and analysis. The thoracic trachea, left lungs and tracheobronchial lymph nodes were fixed in 10% neutral buffered formalin (NBF) and then processed for microscopic examination. The left lung was inflated with 10% NBF to a fixed hydrostatic pressure of 25 cm. After fixation, tissues were trimmed and paraffin embedded for histopathology. The brain and spinal cord were dissected, and sections were reserved for both viral quantification assays (i.e., RT-qPCR and TCID_50_) and histopathology. Samples for RT-qPCR and TCID_50_ were stored at −70 °C or below until processing and analysis. Samples for histopathology were fixed in 10% NBF until trimming, paraffin embedding, sectioning, and staining.

### Histopathology

Formalin-fixed, paraffin-embedded tissues were sectioned and mounted on glass slides. Slides were deparaffinized with xylene and rehydrated with ethyl alcohol and water for staining with hematoxylin and eosin using a Tissue-Tek Automatic Slide Stainer. Microscopic examination was performed by a board-certified veterinary pathologist, and histologic lesions were scored based on the severity and distribution of pathology. Histopathologic changes in the examined tissues were graded semi-quantitatively by a single pathologist on a scale of 0-5. Histopathology was not performed blinded.

### Reverse-Transcription PCR (RT-PCR)

Nasal swabs, CSF, and BALF supernatants were stored at −70 to −90°C until the end of the study, at which time a subset of samples was designated for quantification of EV-D68 genome copies by RT-qPCR. Samples were centrifuged at 4000 x *g* for 5 minutes, and supernatants were harvested. RNA was isolated using the QIAGEN RNeasy Kit or equivalent, according to the manufacturer’s instructions.

EV-D68 genome copies were quantified from extracted RNA samples by RT-qPCR. Genome copies per mL or swab equivalents were calculated from a standard curve generated from RNA standards of known copy concentration. All samples were analyzed in triplicate.

The EV-D68 primers and probe sequences were as follows:

EV-D68 Forward: 5’ CATGAGGCAGTAGCCATTGA 3’

EV-D68 Reverse: 5’ CTGTACTCTCGGTTTCCCATTT 3’

EV-D68 Probe: 6FAM-AACCCACACAACCAGAAACCGCTA-BHQ-1

Amplification and detection was performed using a suitable real-time thermal cycler under the following cycling conditions: 50 °C for 5 minutes, 95 °C for 20 seconds and 40 cycles of 95 °C for 3 seconds, and 60 °C for 30 seconds.

### Median Tissue Culture Infectious Dose (TCID_50_) Assay

Nasal swabs, CSF, BALF supernatants, and terminal tissue samples were shipped to the Pekosz laboratory at Johns Hopkins for processing and determination of infectious viral loads by TCID_50_ assay. In brief, RD cells were plated in 96-well plates, grown to full confluence, and washed with PBS+. Tenfold serial dilutions of the virus inoculum were made, and 200 μL of dilution was added to each of 6 wells in the plate, followed by incubation at 32°C for 5-7 days. Cells were fixed by adding 100 μL of 4% formaldehyde in PBS per well and incubated at room temperature for at least 10 minutes before staining with Naphthol Blue Black solution. TCID_50_ calculations were determined by the Reed-Muench method.

### Western Blot

The antibodies against structural proteins of EV-D68 were detected in a Western blot assay. HeLa cells were infected with a multiplicity of infection (MOI) of 10 of either a Newcastle Disease virus vector co-expressing the precursor of capsid protein P1 and protease 3CD of the EV-D68 (Fermon), or a control vector expressing EGFP and RFP instead of EV-D68 proteins, as previously described (Viktorova et al., 2018). The cells were lysed at 24 hours post-infection, and the lysates were resolved on a 12% or 4-15% gradient PAGE and transferred to a PVDF membrane to generate multiple membrane strips with both lysates located side by side. The 1:1000 dilutions of pre- and post-immunization sera samples from the same animal were incubated with separate membrane strips each containing an EV-D68-positive and a control lane for one hour. After the primary antibody incubation, the membranes were incubated with the goat anti-monkey IgG secondary antibodies conjugated to horseradish peroxidase (ThermoFisher) and developed with Amersham ECL Select chemiluminescent detection reagent (VWR) according to the manufacturers’ protocols. The signal was recorded with an Azure Biosystems C500 digital luminescence detection system. The signals unique to the post-immune sera in the membrane lane containing EV-D68 proteins were considered positive for anti-EV-D68 antibody response.

### Data Analysis

All graphical presentations and statistical analyses were performed using GraphPad Prism software. For RT-qPCR and TCID_50_ assays, samples that were below the lower limit of quantification (LLOQ) for the assay were assigned a value equal to half the LLOQ. Due to the low sample size in each experimental group, no formal statistical analyses were performed as part of this study.

## Acknowledgements

This project was funded in part with Federal funds from the National Institute of Allergy and Infectious Diseases, National Institutes of Health, Department of Health and Human Services, under Contract No. HHSN272201700024I/75N93020F00001/B07. The schematic presented in Figure 1 was prepared using BioRender.com.

## Author Contributions

MSV and AP conceptualized and designed the EV-D68 infection studies. AP and AC propagated and titrated all EV-D68 stocks for animal infections. JD and MSV performed all animal studies. CB and AR performed all RT-qPCR analyses. SMB performed all histopathology. AC and ML performed all TCID_50_ assays. GAB and AZ performed all western blot analyses. JD, DRA and MSV performed all data compilation, analysis and interpretation. All authors participated in the writing of the manuscript and approved the final version.

